# Highly accurate retinotopic maps of the physiological blind spot in human visual cortex

**DOI:** 10.1101/2021.11.29.470502

**Authors:** Poutasi W. B. Urale, Alexander M. Puckett, Ashley York, Derek Arnold, D. Samuel Schwarzkopf

## Abstract

The physiological blind spot is a naturally occurring scotoma corresponding with the optic disc in the retina of each eye. Even during monocular viewing, observers are usually oblivious to the scotoma, in part because the visual system extrapolates information from the surrounding area. Unfortunately, studying this visual field region with neuroimaging has proven difficult, as it occupies only a small part of retinotopic cortex. Here we used functional magnetic resonance imaging and a novel data-driven method for mapping the retinotopic organization in and around the blind spot representation in V1. Our approach allowed for highly accurate reconstructions of the extent of an observer’s blind spot, and out-performed conventional model-based analyses. This method opens exciting opportunities to study the plasticity of receptive fields after visual field loss, and our data add to evidence suggesting that the neural circuitry responsible for impressions of perceptual completion across the physiological blind spot most likely involves regions of extrastriate cortex – beyond V1.

## Introduction

The visual cortex in the occipital lobe contains precise topographic maps, with neighboring locations of the observer’s visual field encoded by adjacent populations of neurons. Damage at various stages of the primary visual pathway results in localized vision loss, such as retinal damage in age-related macular degeneration and glaucoma, as well as stroke or brain trauma that can impact structures at later stages of the human visual hierarchy. While these regions of localized loss of visual sensitivity from damage, known as scotomas, are not entirely irreversible, there is evidence for plasticity by which the brain reorganizes mappings between inputs and neural responses in regions surrounding the region of acquired blindness – either through prolonged experience or active perceptual learning (1, 2). Such plasticity may enable the visual system to compensate for some vision loss, by making better use of intact neural circuitry.

Functional magnetic resonance imaging (fMRI) theoretically enables researchers to study neural processing and map reorganization around the scotomas of human participants in a non-invasive fashion. Advances in neuroimaging technology and data analysis methods have greatly improved the knowledge we can glean about neural functioning in visual cortex. For over a decade, population receptive field (pRF) analysis has become an important method in the toolkit of visual neuroscientists (3, 4). The pRF estimates the position, size, and shape of receptive fields of individual fMRI voxels, corresponding to the aggregate activity of all the neurons within the voxel. However, as a means for better understanding scotomas in patients this comes with several caveats, not least of which is that ground truth is unknown. Functional brain maps of a scotoma patient can only be compared to maps in healthy controls, or possibly to equivalent maps at an intact location of the visual field.

The physiological blind spot is a region of the visual field of each eye that corresponds with the eye’s optic disc. There are no photoreceptors in this region of retina, which causes a naturally occurring scotoma. This seems to make it an ideal model for studying visual scotomas in healthy control participants; however, to date very few investigations of the neural representation of the human blind spot have been attempted. Post-mortem studies of ocular dominance patterns in human area V1 (5) have demonstrated that the physiological blind spot corresponds with a small monocular oval region of V1, subtending a cortical surface area of approximately 50 mm^2^. It is located at an eccentricity of approximately 15°, where cortical magnification is already much reduced compared to the fovea or parafovea (6). This means that even with high spatial resolution brain imaging, only a small number of voxels encode responses from this region of V1. Nonetheless, fMRI experiments have shown that the blind spot is a discontinuity in the visual field representation: with responses to spatially separate stimuli on either side of a blind spot exciting anatomically segregated responses in V1, despite observers perceiving such a configuration as single and continuous (7, 8).

While some important preliminary observations have been made regarding representations of the physiological blind spot in V1, to date no detailed retinotopic maps of this region exist. Here, we report the world’s first detailed mappings of retinotopic organization, in and around representations of the physiological blind spot in V1. Our novel data-driven method will provide researchers with the means for further investigations of this region. More generally, it enables investigations of retinotopic organization around regions of acquired localised blindness which, like the physiological blind spot, are often located in the periphery of human vision – and consequently can pose similar challenges to brain imaging, that we have now solved. Moreover, our data provide converging evidence, suggesting that neural circuits responsible for perceptual completions of form across physiological blind spots most likely have a substrate in extrastriate cortex – beyond V1.

## Materials and Methods

### Participants

Seven observers (ages: 21-41, 3 Female, all right-handed) participated in experiments using a 3 Tesla scanner in the Centre for Advanced MRI at the University of Auckland, New Zealand. Two additional observers (ages: 30 and 37, 1 female, all right-handed) participated in experiments on a 7 Tesla scanner at the Centre for Advanced Imaging at the University of Queensland, Brisbane, Australia. All observers were recruited from staff and student populations at each site, and gave written informed consent to take part. Experimental procedures were approved by the University of Auckland Human Research Participants Ethics Committee, as well as the ethics committee at the University of Queensland.

### Procedure

All participants wore an ophthalmic eye patch over one eye (4 observers: left eye; 3 observers: right eye; both observers in 7 Tesla scans: left eye) before they were placed inside the scanner bore. After initializing the scan, we used a behavioral localizer (see details below) to determine the position and borders of the participant’s physiological blind spot. The center location of the blind spot thus determined was then used to place the stimuli in the retinotopic mapping experiments. Participants underwent 4-6 runs, each lasting 4 mins 10 seconds, for mapping pRFs in the region in and around their blind spot. Following that, the patient bed was moved some ways out of the bore, but the participant remained on it. We moved the eye patch to cover the opposite eye and returned the participant into the bore. The participant then underwent an identical number of runs of pRF mapping, for the same visual field region but now exciting the control eye without a blind spot. Finally, we again moved out the patient bed to remove the eye patch and added the front visor to the 32-channel head coil (at Auckland site only; see scanning parameters). We then acquired a T1-weighted scan of the participant’s brain anatomy. For the first participant at the Brisbane site, we found that the scanner setup restricted their field of view when the control (right) eye was patched. We therefore used the behavioral localizer to map out the visible portion of the stimulus in the control scan for this participant. In the second participant, we instead collected a binocular viewing control (i.e., no eye was patched) to ensure they could see the whole stimulus. All stimuli were generated and displayed in MATLAB using Psychtoolbox 3 (9).

### Blind spot localizer

When the participant had been placed inside the scanner bore, but prior to the functional scans, we localized the physiological blind spot in each observer. The participant fixated a small (diameter: 0.17° of visual angle) black dot located 10.6° of visual angle from the screen center on the side ipsilateral to their patched eye. A radar grid pattern comprising radial lines and concentric circles (10, 11) that emanated from the fixation dot was also displayed to enhance fixation compliance. A larger red disc (diameter: 0.53°) was then moved by the experimenter using the computer mouse. The participant responded verbally via the intercom whenever this red target vanished or reappeared. The experimenter marked this location with a left click on the mouse and the program then placed a small grey dot (diameter: 0.17°) at that location. We moved the target in several directions to ascertain the boundaries of the blind spot, also moving it out of the approximate region to minimize response biases. Occasionally, the experimenter made multiple estimates of the boundary at the same location to improve accuracy. Once the border of the blind spot had been estimated with sufficient detail, a right click on the mouse ended this phase. At this point, geometrically redundant points were removed from the estimated blind spot boundary. The program calculated the centroid location of these border dots and displayed a black dot (diameter: 0.17°) at that location, as well as a line circle (diameter: 10.6°) around it, marking the visual field region to be stimulated during retinotopic mapping. We debriefed the participant to report what they could see. An accurate estimate of the blind spot entails that none of the small dots would be visible, but the large line circle should be clearly visible (meaning it was sufficiently far from the edge of the blind spot). The procedure would be repeated if these criteria were not met although this did not occur.

### Retinotopic mapping stimuli

Retinotopic mapping stimuli were like those used in previous studies (3, 11–13) with a few specific alterations. Participants fixated on a dot aided by the radar grid, identical to the blind spot localizer. Bar stimuli (width: 1.3°) traversed a circular region of the visual field (radius: 10.6°) centered on the blind spot, as determined by the centroid obtained in the localizer (Figure 1A). The bar was only visible within the circular region, so it changed length at each step. One sweep of the bar along a given direction lasted 25 seconds, with one discrete 0.4° step of the bar per second. Each sweep was along the axis perpendicular to the bar orientation. We used eight orientations/directions in sequence starting from horizontal moving upward (0°), and then changing in steps of 45° until we reached the final orientation of 315°. Interspersed between the fourth and fifth sweep and after the eighth sweep, we presented a blank period during which only the fixation dot and radar grid were presented. Inside the bar a checkerboard (side length: 0.95°) flashed with 6Hz.

**Figure 1.**
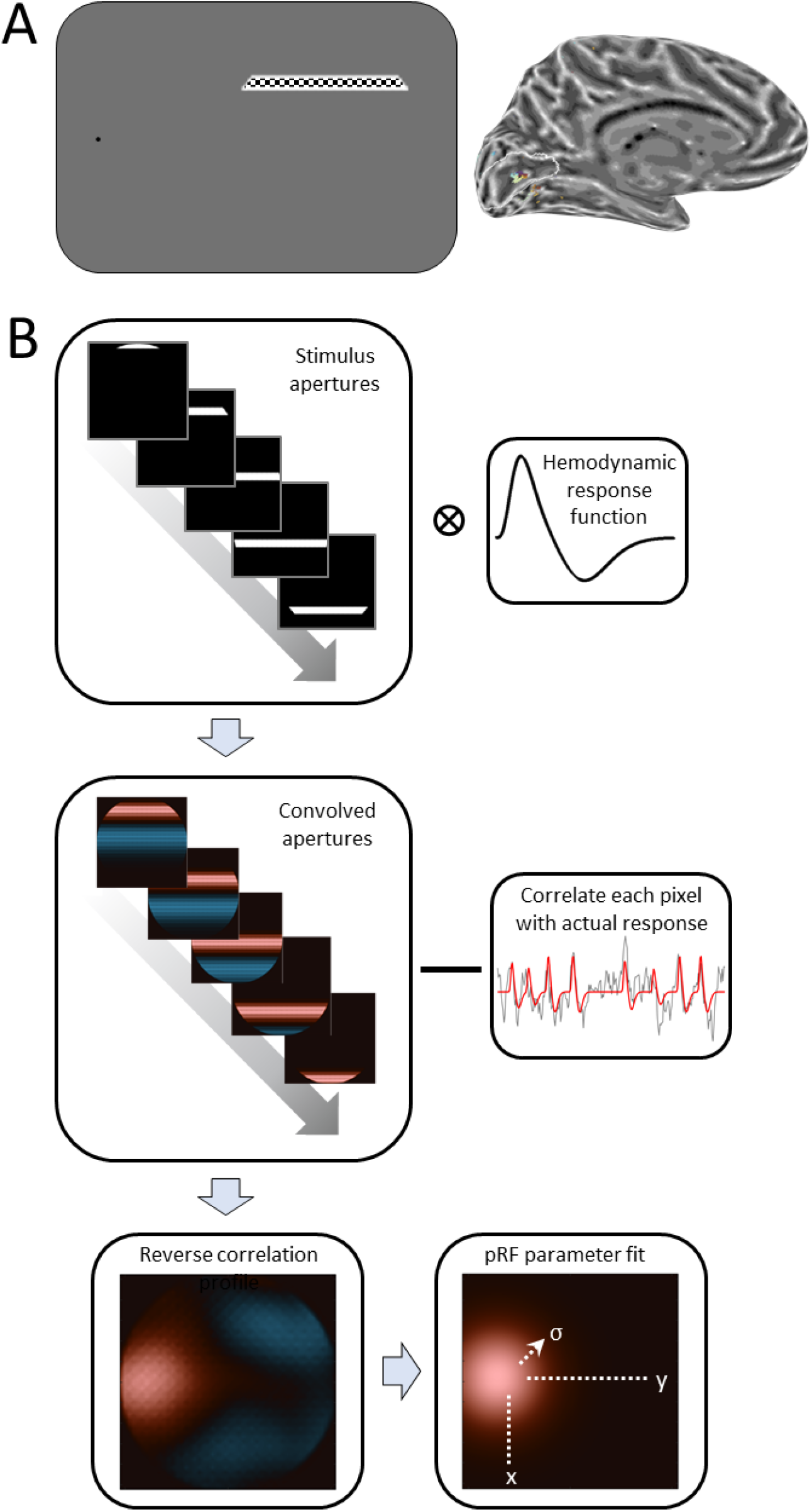
Population receptive field (pRF) mapping at the physiological blind spot. **A**. Stimulus design: Participants monocularly fixated the side of the screen (here: left) while a checkerboard bar stimulus traversed the region of the visual field centered on the blind spot. This drives neural responses in corresponding regions of V1. **B**. Analysis pipeline for reverse correlation pRF fit. Binary apertures indicating the stimulus location relative to the blind spot center at each time point of the experiment are convolved with a canonical hemodynamic response function. To determine the pRF for a given voxel, its actual measured time series is correlated with each pixel time series in convolved apertures. This results in a reverse correlation pRF profile. We then fit a two-dimensional Gaussian pRF model to this profile, to estimate the location (x,y) and size (σ) of the pRF.

To ensure that participants maintained fixation, they performed a detection task. The duration of the scan was divided into brief 200 ms epochs. In each there was a 10% probability that the small black fixation dot would change into a black grapheme. This could be any of the 26 letters of the English alphabet, or a digit from 0-9. Participants were instructed to press a button whenever they saw a number. The 200 ms epoch following a character/digit was always followed by the fixation dot to ensure such events were not too fast.

### Magnetic resonance imaging

At the Auckland site, we used a Siemens SKYRA 3 Tesla scanner with a 32-channel head coil where the front element had been removed to permit an unrestricted view of the screen. This setup results in 20 effective channels covering the back and the sides of the head. Between 4-6 pRF mapping runs of 250 T2*-weighted image volumes were acquired for each eye. We used an accelerated multiband sequence with a TR of 1000 ms and 2.3 mm isotropic voxel resolution, with 36 transverse slices angled to be approximately parallel to the calcarine sulcus. The scan had a TE of 30 ms, flip angle of 62°, field of view 96×96, a multiband/slice acceleration factor of 3, an in-plane/parallel imaging acceleration factor of 2, and rBW was 1680 Hz/Px. After acquiring the functional data, the front portion of the coil was put back on to ensure maximal signal-to-noise levels for collecting a structural scan (a T1-weighted anatomical magnetization-prepared rapid acquisition with gradient echo (MPRAGE) scan with a 1 mm isotropic voxel size and full brain coverage).

At the Brisbane site, data were acquired on a MAGNETOM 7T whole-body research scanner (Siemens Healthcare, Erlangen, Germany) with a 32-channel head coil (Nova Medical, Wilmington, US). B_0_ shimming up to 3rd order was employed to minimize field inhomogeneity. All functional data were collected using the CMRR simultaneous multi-slice (SMS) sequence implementation (https://www.cmrr.umn.edu/multiband) with a matrix size = 128 × 128 × 44 and FOV = 192 × 192 × 66 mm, resulting in an isotropic voxel size of 1.5 mm. The TR was 1466 ms, flip angle = 60°, GRAPPA acceleration factor = 2, SMS acceleration factor = 2, and TE = 30 ms. Whole-brain anatomical images were collected using an MP2RAGE sequence (Marques et al., 2010) with a matrix size of 378 × 420 × 288 and FOV of 201mm x 224mm x 144mm, resulting in an isotropic voxel size of 0.5 mm. The TE was 2.88 ms, TR = 4300 ms, flip angles = 5 and 6°, TI_1_ = 840 ms, and TI_2_ = 2370 ms.

### Data preprocessing

Functional data were realigned and co-registered to the anatomical scan using default parameters in SPM12 (https://www.fil.ion.ucl.ac.uk/spm). We further reconstructed and inflated surface mesh models of the grey-white matter boundary (14–16) using the automatic reconstruction algorithm in FreeSurfer (Version 7.1.1; https://surfer.nmr.mgh.harvard.edu). Functional data were then projected onto this cortical surface model: For each vertex in the surface mesh, we determined the voxel in the functional image that lay at the midpoint between the vertex on the grey-white matter boundary, and the same vertex on the pial surface boundary. We then applied linear detrending to time series for each vertex and run to remove slow drifts, and the time series were normalized to z-scores. We then averaged the runs for the blind spot and control eye, respectively, resulting in two runs of 250 volumes for each condition.

We restricted all further analyses approximately to the occipital lobe by choosing vertices in the inflated surface model whose y-coordinates in FreeSurfer space ≤-35. We calculated a noise ceiling by separately averaging odd and even runs from each condition, and then correlating the time series for each half for each vertex. Using Spearman’s prophesy formula (17) we extrapolated the maximum theoretically achievable correlation for each vertex and squared this value as an estimate of the maximal goodness-of-fit. For expediency, we further limited our analyses to only those vertices with a noise ceiling exceeding 0.15.

### Reverse correlation pRF analysis

In all following analyses, we treated the center of the blind spot as standard pRF mapping analyses would treat the center of gaze. That is, we define that the blind spot center as the center of the mapped visual field. The circular visual field region surrounding the blind spot center had a radius of 5.3°. We created a sequence of stimulus apertures, which depict the location of the bar stimulus for each 1 s fMRI volume with a binary 100×100 pixel matrix, where each pixel denotes whether a stimulus was present at that location, or not (Figure 1B, top). To account for the lag in the blood oxygenation dependent response, we convolved the sequence of stimulus apertures for bar positions, across the 250 s run, with the canonical hemodynamic response function based on previously collected data (18). We then used these convolved apertures for a reverse correlation analysis (see Figure 1B, middle). Separately for each vertex of the occipital lobe, we calculated a linear regression of the observed fMRI time series using the time series for each pixel of stimulus apertures as regressors. We also included a global covariate, to estimate the intercept. This generated a pRF profile for that vertex showing how strongly each visual field location had correlated with the observed fMRI response (Figure 1B, bottom-left). We estimated the pRF location by determining the profile maximum. We also recorded the squared correlation R^2^_RC_ between this pixel and the fMRI response.

Resulting pRF profiles can be asymmetric, or have multiple peaks. Given our use of a regular bar stimulus, profiles can also contain artifacts related to spatiotemporal correlations within the stimulus sequence. Consequently, we restricted further analyses to vertices where R^2^_RC_>0.1. We then fit a two-dimensional pRF model to these reverse correlation profiles, to estimate the location and extent of pRFs. We used a symmetric two-dimensional Gaussian profile (Figure 1B, bottom-right), defined by its x and y positions, its standard deviation σ, and amplitude β. The goodness of pRF fit (R^2^_pRF_) is determined by calculating the sum of squared pixel-wise residuals between the predicted pRF profile and the one estimated from reverse correlation. All further quantitative analyses of data are restricted to vertices where R^2^_pRF_>0.5.

### Forward-model pRF analysis

We also conducted a standard pRF analysis using forward-modelling (3, 11, 19, 20). This analysis fits a predicted time series of the fMRI response to the observed time series, based on known stimulus apertures and an assumed pRF model profile. In short, a neural response prediction is generated by overlaying the stimulus aperture onto a two-dimensional Gaussian pRF model. This predicted response time series is further convolved with the canonical hemodynamic response. The best fitting pRF model is determined by varying its visual field position (x,y) and size (σ) parameters, and calculating the correlation between the predicted and observed time series. This fitting uses a coarse-to-fine procedure: first, we generate several thousand predictions for a plausible range of permutations of the three pRF parameters, and calculate an extensive grid search by finding the best-correlated combination. These parameters are then used to seed an optimization procedure (21, 22) to fine-tune the fit. Because correlation is scale-invariant, we also calculated a linear regression between this fit and the observed time series, to determine the amplitude (β_1_) and baseline level (β_0_) of the response. These pRF modelling procedures have been described in detail elsewhere (11, 19). However, here we used a different biophysical model for predicting a pRF’s neural response than in previous work by us and others. Specifically, we calculated the percentage of overlap between the pRF model profile and the stimulus aperture. This accounts for the fact that the response to a constant visual stimulus should vary with different pRF sizes. Note that this merely affects the modelled signal amplitude, not the pRF position or shape.

Our main analysis used the stimulus apertures of the full bar stimuli as they would have appeared on the screen (and which were used for the reverse correlation analysis). Previous research has shown, however, that pRF estimates derived via forward-modelling are susceptible to potential biases when a scotoma is not explicitly modelled in the stimulus apertures (23). This could produce spurious estimates of pRFs inside the blind region. We therefore generated masked stimulus apertures, specific for each individual participant, based on the blind spot localizer. Within the polygon described by the blind spot border, the aperture was set to zero. This mask was smoothed with a Gaussian kernel of 0.4°, to make the edges of the scotoma more biologically plausible. We then repeated the forward-model pRF analysis using this masked aperture.

### Regions of interest

All quantitative analyses of the blind spot were restricted to a probabilistic estimate of V1, based on an anatomical atlas derived from postmortem brains (24). We selected a continuous region of this atlas, predicted to be within V1 with ≥80%. In all participants this revealed clusters of significant pRF fits (especially for the reverse correlation approach) at a location consistent with the expected representation of the blind spot in V1 (5, 7). However, due to artifacts a few stray vertices occasionally survived thresholding throughout the rest of V1. We therefore further defined a circular region of interest (ROI) around the main cluster. Using the map for the control condition, we manually selected the vertex approximately at the center of the cluster and then used a geodesic region growing procedure by taking 12 steps across the grey-white matter surface mesh away from that center. This ROI encompassed the significant cluster. All further analyses reported were conducted exclusively on vertices from this ROI.

## Results

### Retinotopic maps around the blind spot

Figure 2A shows maps of the angular (radial) coordinates of pRFs relative to the blind spot center for one participant. In maps for both the control and blind spot scans, a cluster of significant retinotopic responses is visible at the location where the blind spot is expected to be encoded, approximately two thirds along the calcarine sulcus within V1 (5). Further, the estimated radial position progressed smoothly around the entire color pinwheel, indicating that neighboring vertices of the surface model encoded pRFs adjacent in the visual field. The radial location of pRFs also corresponded well with the expected location based on the retinotopic organization of V1, such that more foveal (light blue-yellow-light orange) and peripheral (orange-purple-blue) sides of the blind spot corresponded with vertices facing the posterior and anterior sides of V1, respectively. Similarly, the inferior and superior vertices of the cluster encoded upper (orange) and lower (blue) visual field locations. In some participants, clusters were also visible outside of V1, at what is presumably the border between V2 and V3. On the ventral side, Figure 2A shows a cluster in the lingual gyrus, which predominantly encoded upper visual field locations (yellow-orange-purple) consistent with a reversal across the horizontal meridian, which intersects with the blind spot. An equivalent cluster encoding lower visual field locations (blue) can be seen on the dorsal side in the cuneus.

**Figure 2.**
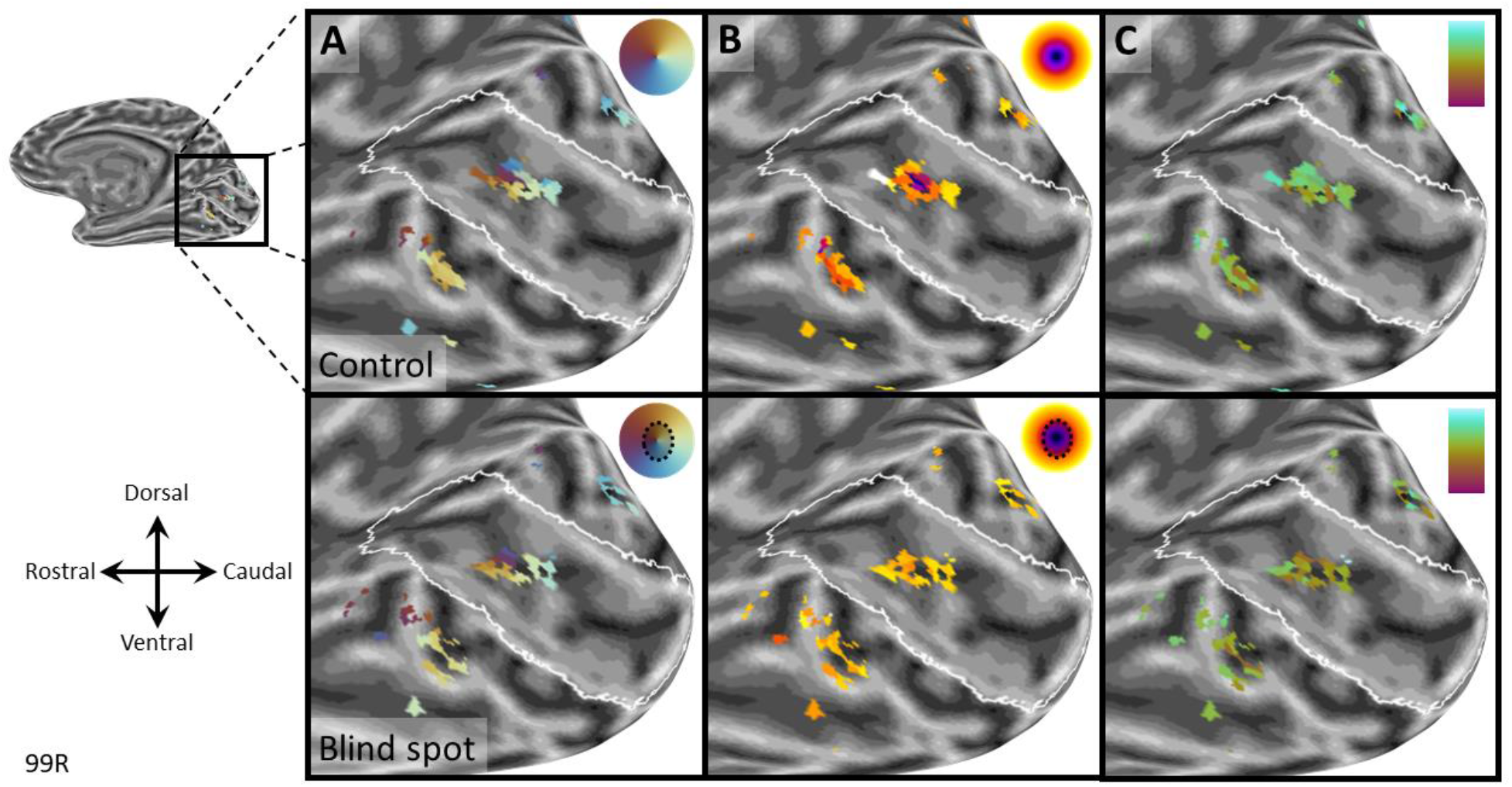
Retinotopic maps form one observer scanned at 3 Tesla. Parameter estimates from pRF model fit (R^2^_pRF_>0.5) are shown on an inflated model of the right occipital lobe. *Top row:* Control map with left eye patched. *Bottom row:* Blind spot map with right eye patched. **A**. Radial pRF position relative to blind spot center. **B**. Distance from blind spot center (color scale: 0-6°). **C**. pRF size (color scale: 1_magenta_-3_cyan_°). The white outline is a prediction of the borders of V1 based on cortical folding (24).

Maps for distance from the blind spot center were less clear. Nevertheless, for the control eye, a concentric organization is visible (Figure 2B, top). In the blind spot condition, not much difference could be determined between vertices (Figure 2B, bottom). This is unsurprising, however, as the blind spot effectively removes central vertices from these clusters. Notwithstanding this, differences between blind spot and control maps reveals a more complete coverage of the mapped region in the control condition. Taken together, these maps demonstrate that our experiments activated the expected clusters of V1 vertices, and that despite their small size, these clusters contained retinotopic maps consistent with the corresponding visual field location.

To obtain more detailed maps, we carried out similar experiments in two participants at 7 Tesla using a finer voxel resolution (1.5 mm isotropic voxels). Indeed, results showed an even clearer pinwheel progression of radial coordinates (Figure 3A), and a distinct concentric gradient in pRF distances from the blind spot center in control scans (Figure 3B, top). The blind spot condition resulted in a doughnut shaped cluster, containing the same pinwheel organization of radial coordinates and pRFs on the outer edge of the mapped portion of the visual field (Figure 3A-B, bottom). In addition, our estimate of pRF size showed a gradient of increasing sizes (Figure 3C; brown-orange-green gradient) for vertices from the foveal (anatomically caudal) toward the peripheral (anatomically rostral) side. This is consistent with the well-established relationship of pRF sizes increasing with eccentricity (3, 6).

**Figure 3.**
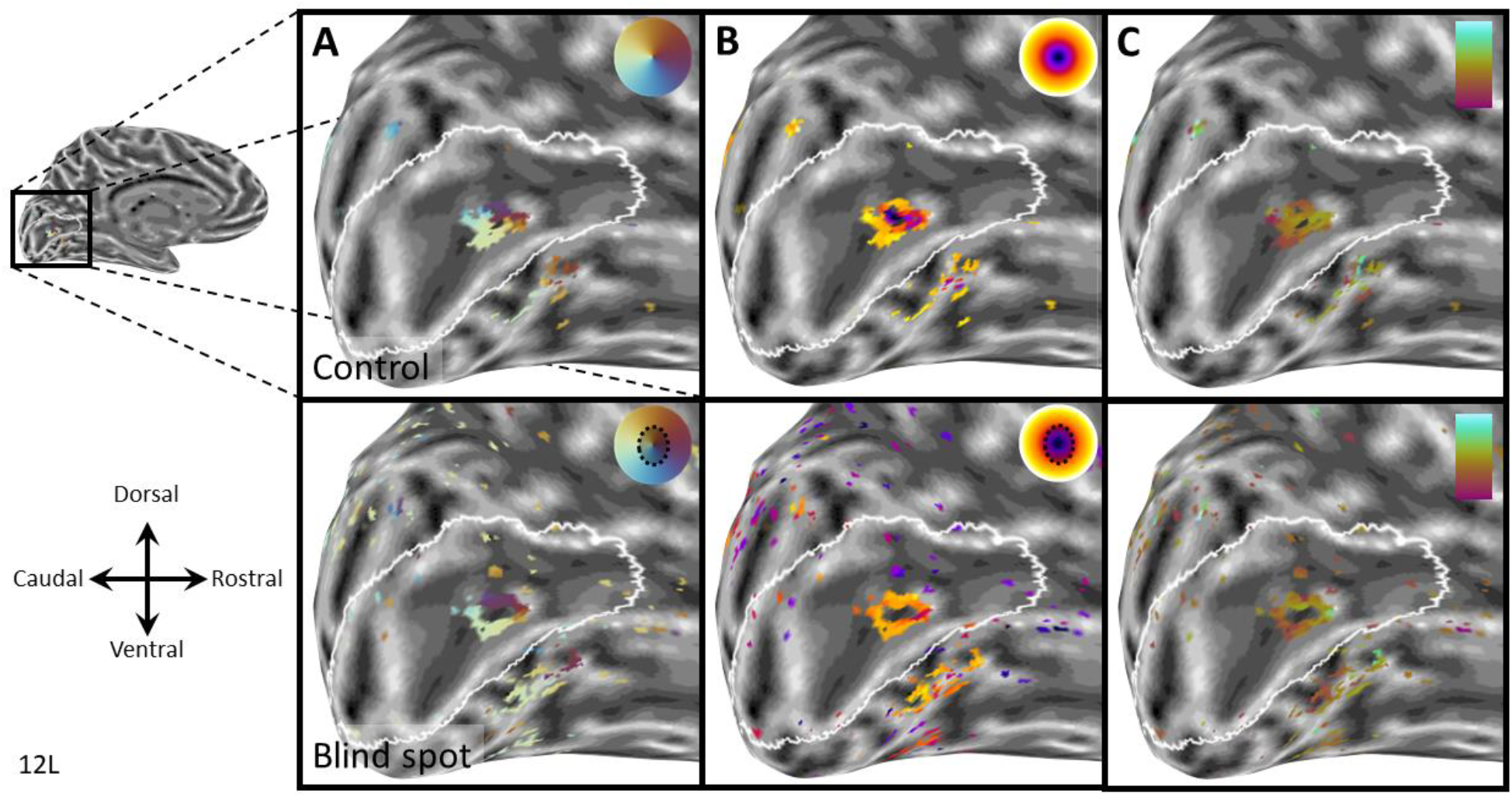
Retinotopic maps form one observer at 7 Tesla. Parameter estimates from pRF model fit (R^2^_pRF_>0.5) are shown on an inflated model of the right occipital lobe. *Top row:* Control map with binocular viewing. *Bottom row:* Blind spot map with left eye patched. **A**. Radial pRF position relative to blind spot center. **B**. Distance from blind spot center (color scale: 0-6°). **C**. pRF size (color scale: 1_magenta_-3_cyan_°). The white outline is a prediction of the borders of V1 based on cortical folding (24).

Next, we quantified the gradient of pRF size when moving from the foveal to the peripheral side of the blind spot. We selected all vertices with R^2^_pRF_>0.5 whose horizontal visual field position was within 6° of the blind spot center. Separately for each participant, and each condition, we fit a robust linear regression to estimate the slope of the gradient. To make data comparable across participants whose left and right hemispheres were scanned, we inverted the sign for horizontal pRF positions for the right hemisphere. Thus, negative horizontal positions correspond to the foveal side and positive positions to the peripheral side. This revealed consistent positive slopes in all participants (Figure 4), both for the control eye (one-sample t-test vs 0: t(8)=6.26, p=0.00024) and the blind spot condition (t(8)=4.01, p=0.00391). Average slopes for the two conditions were also not significantly different (t(8)=1.56, p=0.15763). This demonstrates that in both conditions, pRF size increased from more central to peripheral locations, even when that gradient was not always obvious in all cortical maps obtained at 3 Tesla.

**Figure 4.**
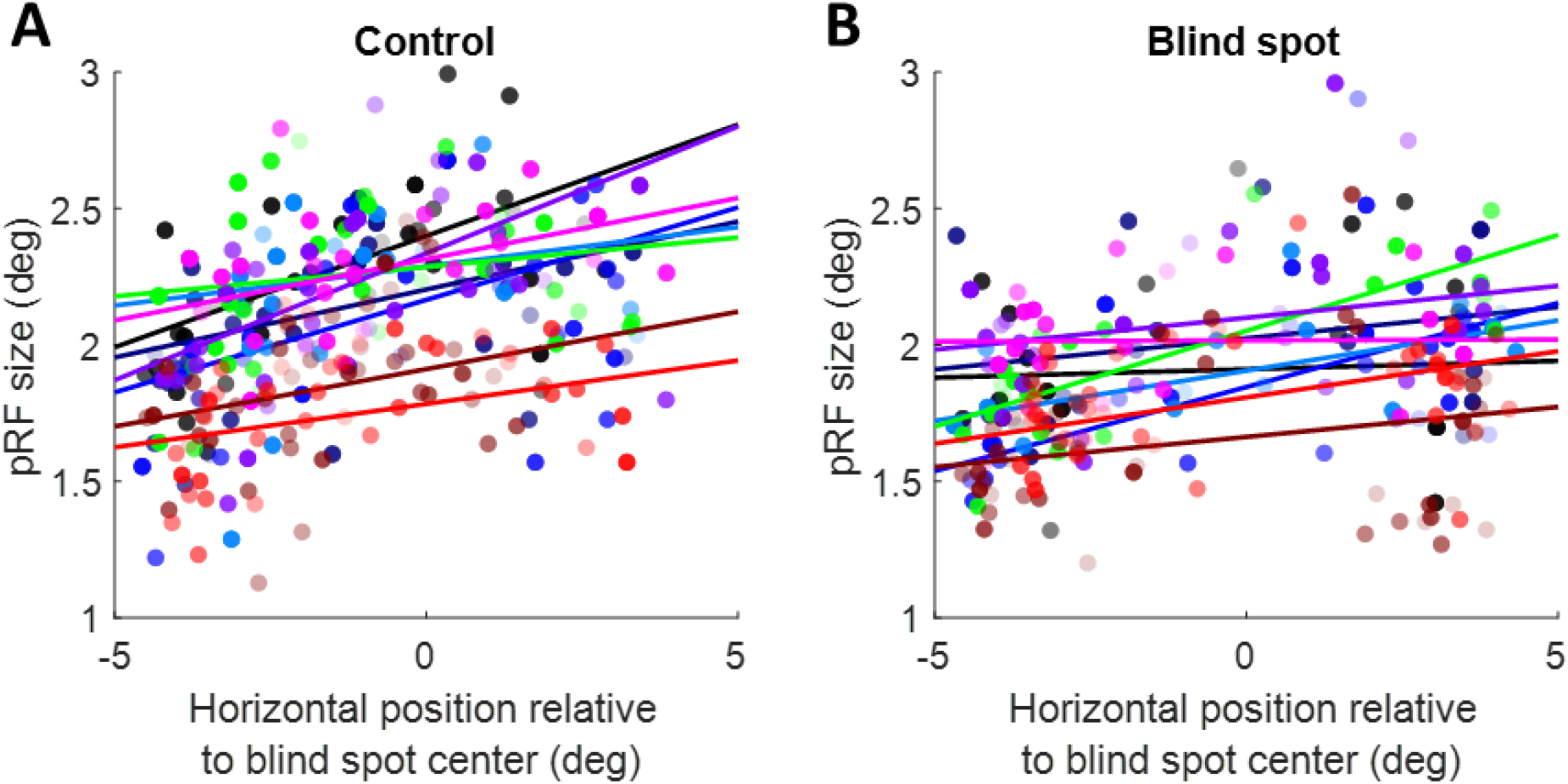
Quantifying pRF size gradient across the visual field. pRF sizes from the control eye stimulation (A) or blind spot eye stimulation (B) were plotted against the horizontal pRF location relative to the blind spot center (negative and positive values, respectively, denote visual field locations on the foveal side and peripheral side of the blind spot). Each dot denotes parameters from one pRF. Solid lines indicate a robust linear regression fit. Colors indicate the 9 different participants.

### Visual field coverage of reverse correlation profiles

We next used the reverse correlation profiles of pRFs to reconstruct how much of the visual field was covered by these pRFs. Peak amplitudes of the profiles from all significant vertices were normalized to their maximum, and all values below 50% of the maximum were set to zero. This effectively restricts the pRF profile to the full-width-at-half maximum. We then averaged these restricted profiles and plotted visual field coverage. In the control condition, the whole mapped region of the visual field is covered by pRFs, albeit not homogeneously (Figure 5A). In particular, the density of pRFs was greater on the side facing the fovea than the periphery, consistent with the gradient of cortical magnification and receptive field density in V1 (6). We obtained similar maps for the blind spot condition (Figure 5B), but notably these receptive field profiles spared the locations of the blind spot scotoma (as determined by the behavioral localizer) in all participants. Due to the inhomogeneity in coverage, the annulus regions around the blind spot contains some gaps. Nevertheless, this suggests the reverse correlation profiles precisely captured the extent of blindness.

**Figure 5.**
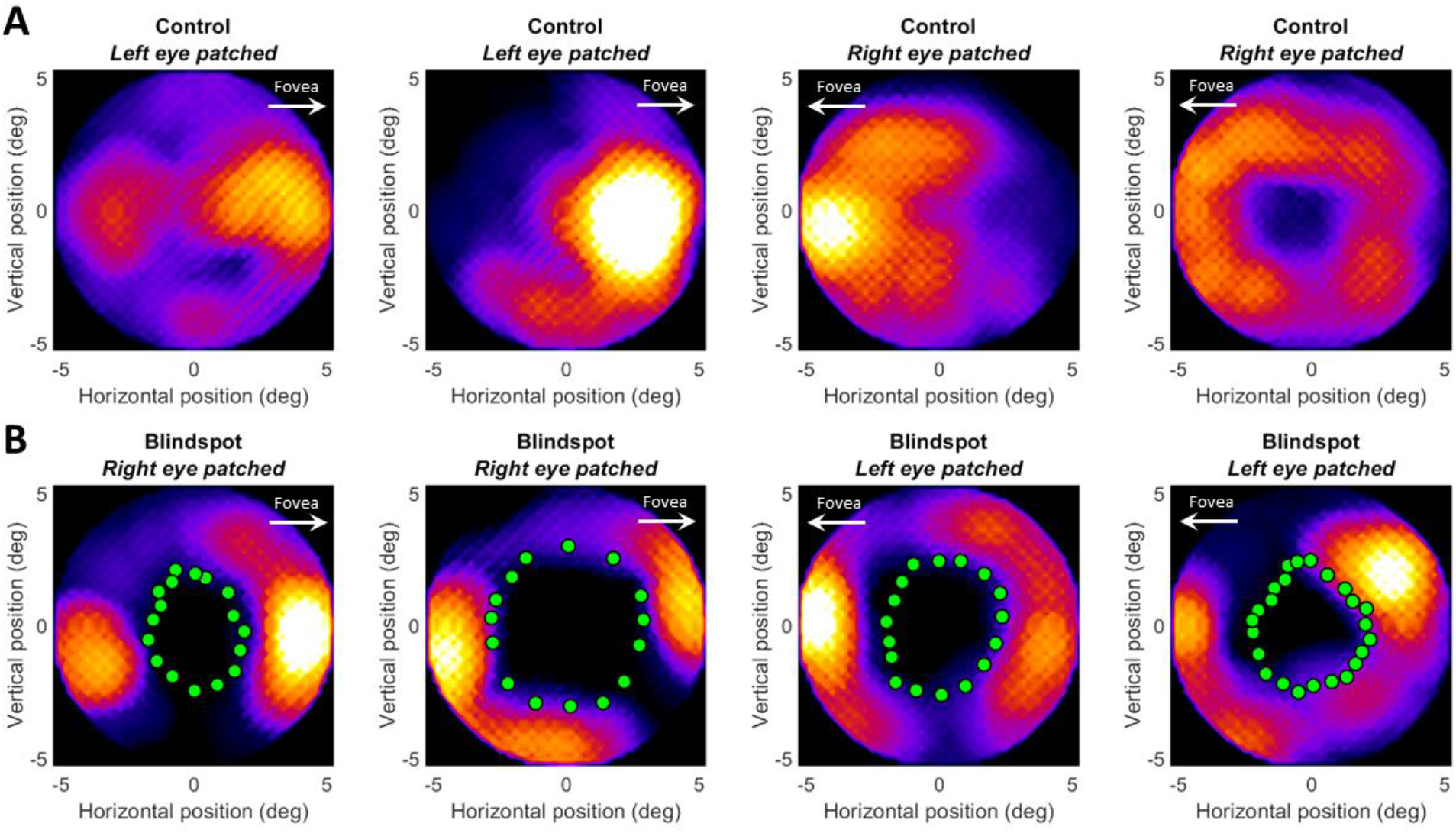
Reconstruction of retinotopic maps in visual space based on reverse correlation profiles. The heat map indicates the density of pRF profiles at a given visual field location. Coordinates are in degrees of visual angle relative to the blind spot center. Data from four participants are shown in columns. **A**. Control eye stimulation. **B**. Blind spot stimulation. Green dots denote the outline of the blind spot as determined by the behavioral localizer.

### Visual field reconstruction of pRF fits

To visualize the position of pRFs directly, we next created scatter plots of pRF position and size (defined by a radius of 1σ) in visual space (Figure 6). Consistent with the visual field coverage (Figure 5) this showed complete coverage of the mapped visual field region in the control condition (Figure 6A). In the blind spot condition, pRF centers spared the blind spot scotoma (Figure 6B), although many Gaussian pRF fits extend into the scotoma. As suggested by the reverse correlation profiles, the density of pRFs in the annulus region surrounding the blind spot varied. Nevertheless, the entire region outside the scotoma was covered by pRFs.

**Figure 6.**
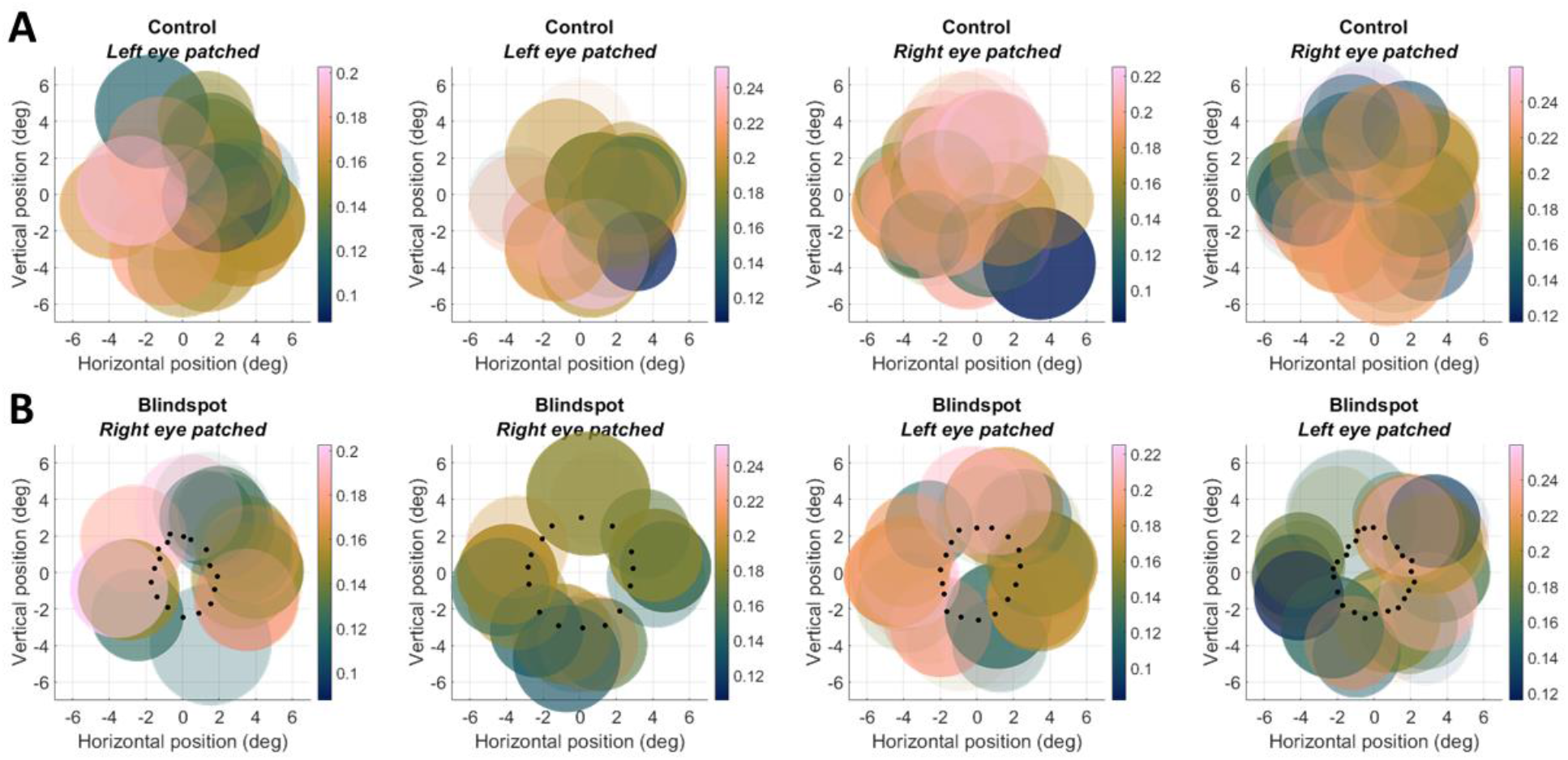
pRF model fits to reverse correlation profiles. The scatter plots denote the location and size of pRFs. The color scale indicates the peak amplitude. Coordinates are in degrees of visual angle relative to the blind spot center. Data from the same four participants as in Figure 5 are shown in columns. A. Control eye stimulation. B. Blind spot stimulation. Black dots denote the outline of the blind spot as determined by the behavioral localizer. the outline of the blind spot as determined by the behavioral localizer.

To quantify the visual field coverage, we divided pRFs from the control condition into those whose centers fell inside the scotoma, and those that fell outside of it. Figure 7 plots the mean peak amplitudes for pRFs outside the scotoma against pRFs inside the scotoma for each participant. Data from the control condition (blue circles) clustered around the identity line with half of data points above and half below it. This shows that responses were comparable for pRFs inside and outside the scotoma. In contrast, data from the blind spot condition suggested weaker responses for pRFs inside the scotoma than outside it; only one participant had a data point above the identity line. It is, however, important to stress that using the control condition to define this region of interest incorporates not only the true pRF position but also any error inherent in mapping the blind spot’s location and extent. Some pRFs estimated as falling inside the scotoma might truly be located outside of it, and vice versa. Hence, this analysis necessarily underestimates differences between these regions.

**Figure 7.**
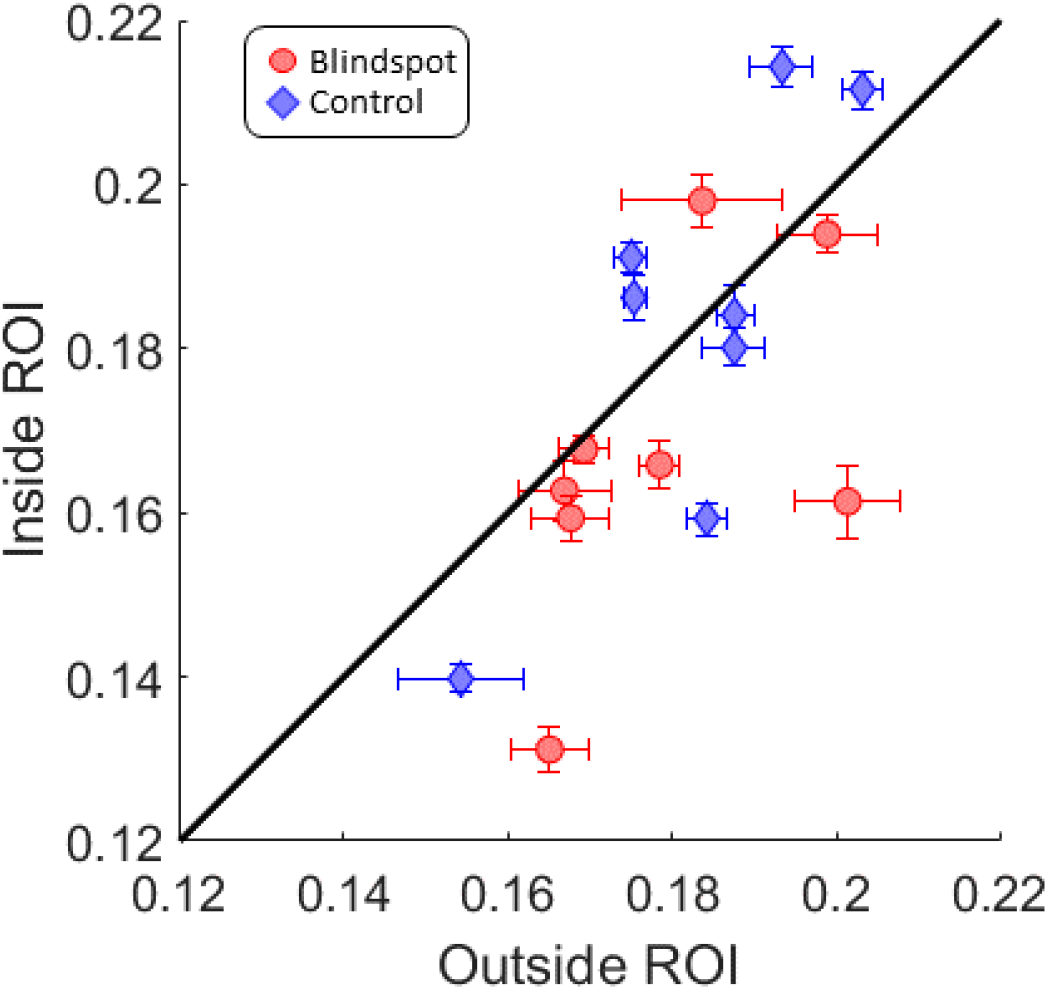
Mean V1 response per participant in and around the blind spot. The mean response amplitude of pRFs in the blind spot region of interest (ROI) plotted against the mean of pRFs outside it, based on the control map. Each symbol denotes one participant. Error bars denote the standard error of the mean across pRFs. Blue diamonds: Control map. Red circles: Blind spot map.

Please note that data from the first participant scanned at 7 Tesla are not included in this plot, as the incomplete field of view in the control condition (when the opposite control eye was patched) precluded us from carrying out this analysis. This incomplete field of view, however, afforded us with a serendipitous opportunity to further test the accuracy of this approach for mapping visual field scotomas. For this participant we show the visual field coverage and pRF scatter plot separately in Figure 8. As with other participants, the mapping approach allowed precise localization of significantly activated pRFs sparing the blind spot scotoma (Figure 8C-D). Critically, in the control condition, these reconstructions also distinctly respected the boundary of the visible portion of the visual field (see green/black dots in Figure 8A-B). This demonstrates that the approach could also map the broader absence of stimulation in these scans. The method should generalize to other kinds of visual field loss than that due to the physiological blind spot.

**Figure 8.**
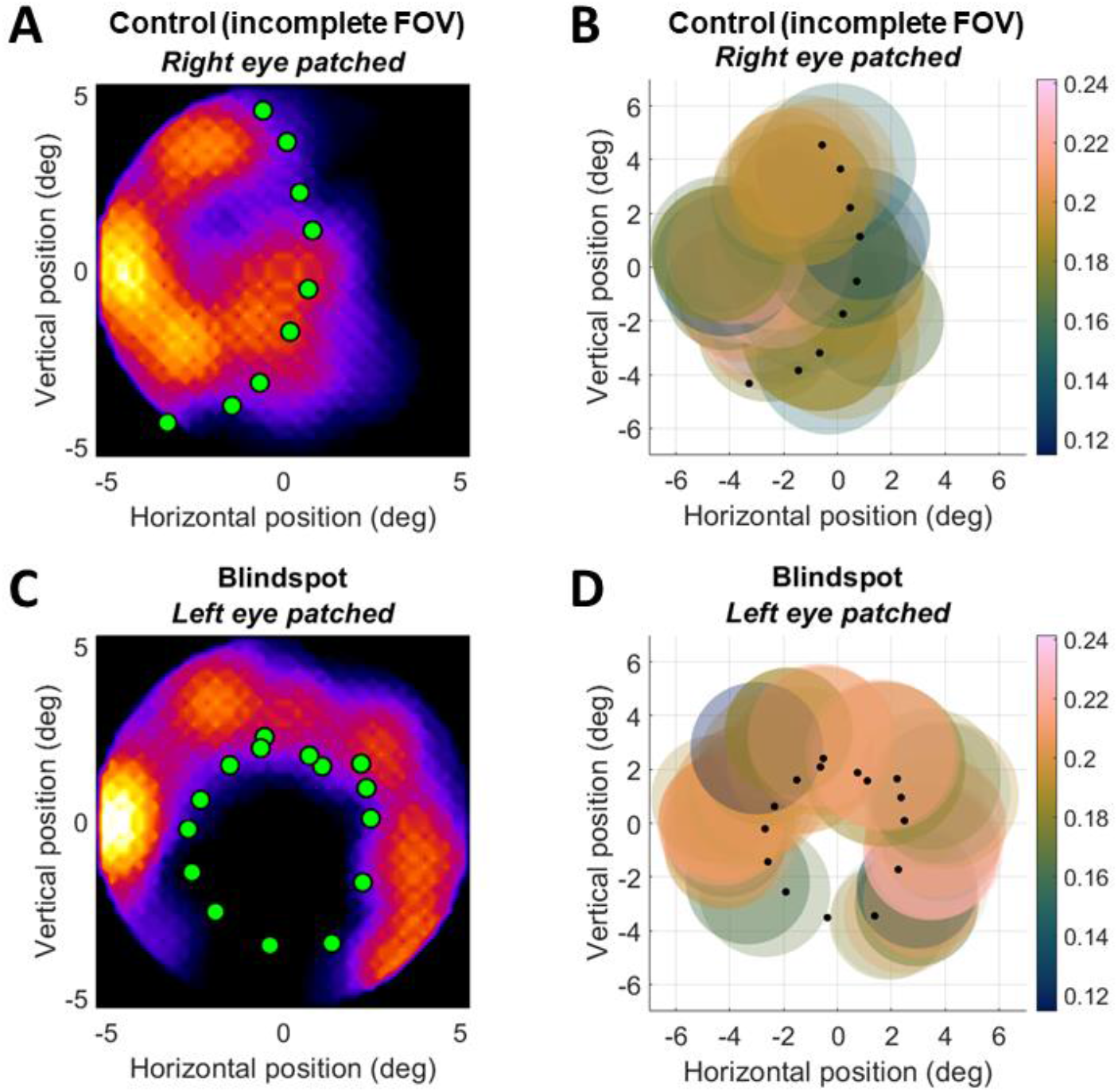
Reconstruction in visual space (**A**,**C**) and corresponding pRF model fits to reverse correlation profiles (**B**,**D**). The scatter plots denote the location and size of pRFs. All conventions as in Figures 4 and 5, respectively. Data is from a participant scanned at 7 Tesla who had an incomplete field of view (FOV) with their control eye. **A-B**. Control eye stimulation. **C-D**. Blind spot stimulation. Green/black dots denote the visible field of view (**A**,**B**) or the outline of the blind spot (**C**,**D**) as determined by the behavioral localizer.

### Comparison with forward-model pRF fits

We also analyzed our data using a more traditional pRF analysis based on forward-modelling (3, 10, 11). Figure 9 shows scatter plots of pRFs analyzed using this approach, comparable to Figure 6. As with our reverse correlation data, in the control condition pRFs covered most of the mapped visual field region, albeit less completely (Figure 9A). Moreover, in the blind spot condition, pRF centers exclusively spared the blind spot scotoma (Figure 9B). However, only a small number of vertices survived statistical thresholding, so these maps are generally sparse and incomplete. The standard forward-modelling approach was evidently less effective in estimating the extent of the scotoma.

**Figure 9.**
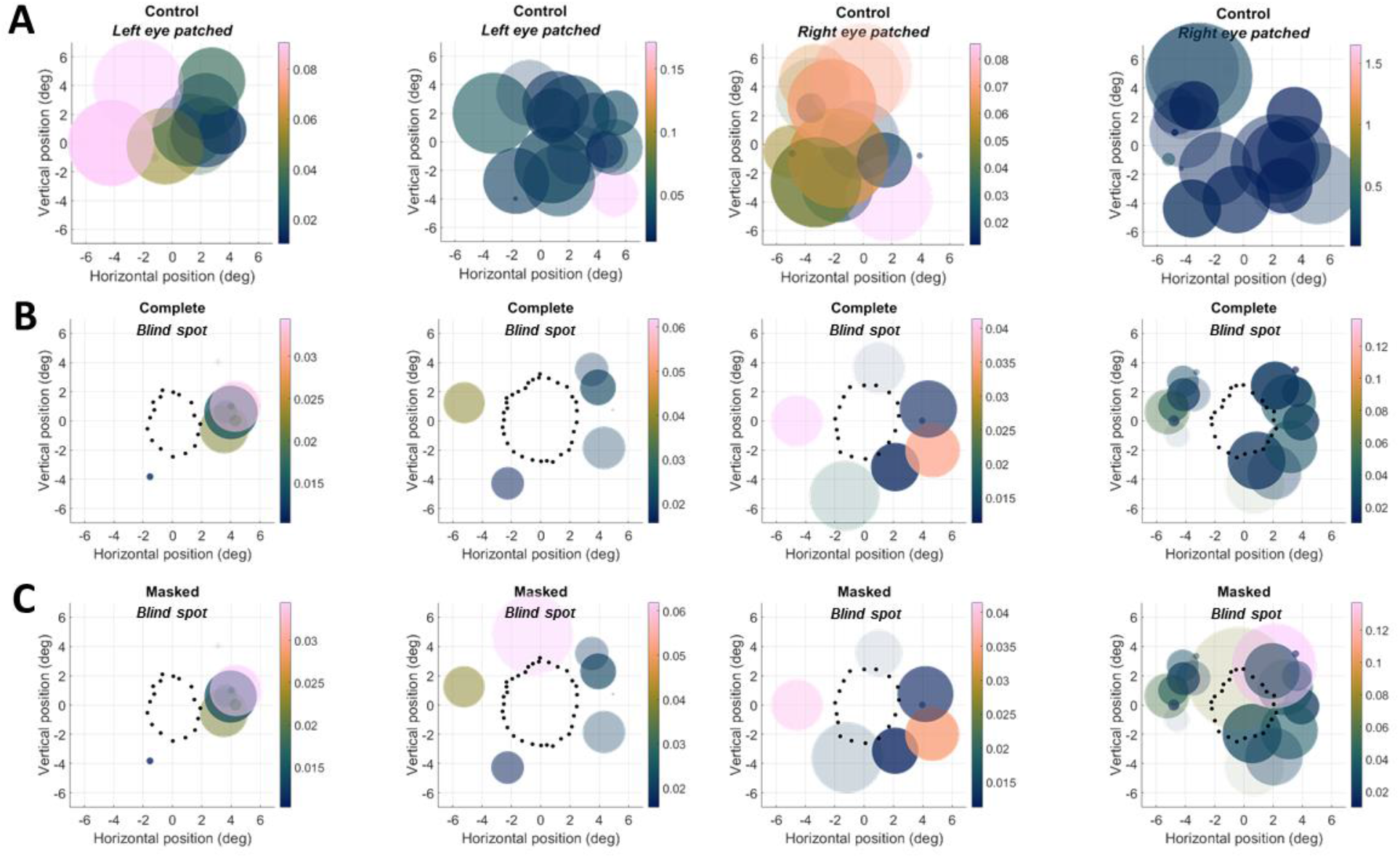
pRF model fits using forward-model analysis. The scatter plots denote the location and size of pRFs. The color scale indicates the response amplitude (β). Coordinates are in degrees of visual angle relative to the blind spot center. Data from the same four participants as in Figure 5 are shown in columns. **A**. Control eye stimulation. **B**. Blind spot stimulation. **C**. Blind spot stimulation using masked apertures to account for scotoma. In B-C, black dots denote the outline of the blind spot as determined by the behavioral localizer.

To further quantify the reliability of pRF positions for the two analysis methods, we reasoned that for pRFs located outside the blind spot the position estimates should be similar for the control and blind spot condition. We therefore selected the pRFs that survived statistical thresholding in both conditions, and then computed the mean Euclidian distance between the blind spot and control estimates for these pRFs, separately for the reverse correlation and the forward-model analysis. This analysis revealed significantly smaller distances in pRF positions (t(7)=-3.6, p=0.0087) for the reverse correlation (mean=1.66°) than the forward-model (mean=2.59°).

Previous research suggested that when using standard pRF modelling techniques for mapping scotomas it is important that the stimulus model incorporates the extent of the scotoma (23). Without this, the modelling approach can produce biased estimates of pRF position. Most notably, this should result in pRFs being falsely estimated to fall inside the scotoma, something we did not see in our data at all (Figure 9B). We nevertheless conducted a control analysis, using stimulus apertures for each participant in which the blind spot scotoma had been masked out. The results for these pRF maps were extremely similar to those obtained with the complete stimulus apertures (Figure 9C). This demonstrates that the poor quality of forward-model pRF estimates in the blind spot condition were not simply due to using an incorrect stimulus model.

## Discussion

We used pRF analysis based on reverse correlation to map the retinotopic organization of the physiological blind spot in human observers. We obtained highly accurate maps of the visual space surrounding the blind spot. In particular, the organization of radial (angular) pRF position relative to the blind spot center followed the pattern expected based on the known retinotopic organization of V1. Maps for the distance from the blind spot center, and especially pRF size, were less clear in participants scanned at 3 Tesla. This is likely due to the comparably coarse voxel resolution relative to the size of the blind spot. Two participants scanned at 7 Tesla, with 1.5 mm isotropic voxel resolution, confirmed expected map gradients for these pRF parameters. We further projected pRFs back into visual space to plot the visual field coverage of these small retinotopic maps. This showed accurate delineation of brain activity to only those sub-regions of the visual field outside the scotoma. In contrast, a conventional forward-modelling approach for estimating pRFs was far less accurate. While pRFs estimated in that way also consistently spared the blind spot scotoma, they were sparse and afforded incomplete visual field coverage. Control analyses confirmed this was not trivially explained by failing to account for the scotoma in the pRF model (23).

The study of retinotopic maps and reorganization around regions of vision loss is hampered by several methodological limitations. Testing and validating the techniques for visual field mapping requires a well-defined model of how the scotoma should behave. Some researchers have therefore used masks to block out pre-defined parts of the visual field to validate measurements (23). These efforts are important, and have resulted in several notable insights – for example demonstrating that estimates of preferred locations near scotomas can be biased, especially when the scotoma is not factored into the analysis (23). Yet masking the stimulus by replacing it with a blank region on the screen is not the same as an actual scotoma. This approach can therefore only inform about analytical issues, and not about any physiological peculiarities regarding the consequences of visual stimulation within and immediately surrounding the scotoma.

Other studies have investigated cortical representations of the foveal rod scotoma (25, 26), which arises because macula contains predominantly cone photoreceptors. However, such experiments require scotopic stimulation (ideally after dark adaptation), rendering them difficult to apply broadly and limiting the generality of these results.

Another recent study used an elegant design to map artificial scotomas – a visual illusion in which a dynamically flickering background fills in a static region of the stimulus, effectively simulating temporary blindness within that region (27). While fascinating, this is probably not directly comparable to an actual scotoma, arising due to an absence of innervation/stimulation. Differences in terms of physiological processes might be considerable (28, 29) – notwithstanding the conceptual interest of that study. In contrast, the physiological blind spot is a visual field region corresponding to the optic disc where there are no retinal photoreceptors. This makes it a perfect model for a small, localized scotoma, as might appear after retinal damage or lesions to primary visual cortex.

Our results demonstrate that even at relatively conventional voxel resolutions, it is possible to produce accurate maps of visual scotomas. This opens a promising avenue for studying map reorganization and plasticity after damage to the visual pathway (1). Previous research has found no evidence for large-scale changes in retinotopic maps after macular degeneration (30). However, previous experiments might have been susceptible to limitations of previous analysis procedures (23), or plasticity effects might have been too subtle to be resolved by those experiments. The precision with which our reverse correlation maps could delineate the borders of the blind spot scotoma suggests it could be a more powerful tool for exploring small-scale changes.

Our method also opens opportunities for studying active perceptual processing. Under normal binocular viewing conditions, input from the fellow eye compensates for the region of blindness resulting from the physiological blind spot, and even with monocular viewing the visual system can fill in missing information by perceptually extrapolating patterns from the regions directly abutting the blind spot. Maps obtained using our method could be used to study the neural correlates of this perceptual completion, at a level of detail thus far reserved for invasive electrophysiological recordings in animal models (31–33).

Further refinements to our method could be considered. Using reverse correlation of mapped pRFs is not new. Rather, this technique has been used in a number of studies (18, 34–36). Some of these used analysis approaches (e.g. using ridge regression) that account for the considerable spatiotemporal correlation in conventional pRF stimulation paradigms (12, 37). We decided against this, as these modelling approaches drastically increase processing time. Despite spatiotemporal correlations in the stimulus sequence, our approach evidently produced precise maps of the blind spot scotoma. However, future studies might consider using such refinements, e.g. when studying subtle changes in pRF parameters around the blind spot. Randomized stimulation sequences that break spatiotemporal correlations might further enhance the accuracy of these maps, although they are also associated with poorer signal-to-noise ratios (23, 37, 38).

While we made no attempt to quantify the effect, our traversing bar stimulus induced at least some perceptual completion, in that none of our participants reported that the moving bars used in mapping seemed to split in two as they passed through the blind spot. It is therefore notable that no responses were observed in regions of V1 corresponding with the site of the physiological blind spot. This is consistent with previous fMRI experiments (7), which similarly found no representation of the physiological blind spot in V1. The convergence of this evidence suggests that perceptual filling-in likely arises through circuits involving extrastriate brain regions – beyond V1.

## Acknowledgements

We thank Roberta Gough and Lydia Zhu for help with data collection. We further thank the team at the Center for Advanced MRI and Nicole Atcheson and Aiman Al-Najjar (Centre for Advanced Imaging) for their invaluable support.

## Code and results availability

The stimulus presentation code is available at https://osf.io/m6tqn. This repository also contains data plots for all participants in this study which could not be shown here. Local data protection rules and our ethical approval do not allow us to share native space brain data publicly. Data may be shared conditionally by the corresponding author upon request.

## Competing interests

The authors declare there are no competing financial or non-financial interests.

## Notes

https://osf.io/m6tqn

### Competing Interest Statement

The authors have declared no competing interest.

